# AI-Driven Discovery and BSL-4 Validation of Cross-Filovirus Ebola-Marburg Inhibitors and their Synergistic Combinations

**DOI:** 10.64898/2026.07.09.737586

**Authors:** Holli-Joi Martin, Marcus Tullius Scotti, Sankalp Jain, Laura K. McMullan, Payel Chatterjee, Cleber Melo-Filo, Maximilian Caza, Alexander Tropsha, Hsiu-Ling Lin, Michael Flint, Emily Lee, Michael K. Lo, Alexey V. Zakharov, Eugene N. Muratov

## Abstract

Filovirus outbreaks caused by Ebola virus (EBOV) and Marburg virus (MARV), pose severe global health threats characterized by high rates of fatal hemorrhagic fever. While species-specific vaccines and therapeutic monoclonal antibodies are approved for Zaire ebolavirus, broadly-active therapeutics remain unavailable, leaving populations vulnerable to MARV and other pathogenic Ebola species, such as Bundibugyo (BDBV) and Sudan (SUDV) ebolaviruses. Here we report a computationally guided, infectious virus validated screening platform for the rapid discovery of broad-spectrum filovirus antivirals. By leveraging quantitative structure–activity relationship (QSAR) models, we screened 142,382 compounds *in silico* to prioritize 125 high-potential candidates. Subsequent dose–response and viability profiling identified 23 compounds exhibiting potent, low-micromolar pan-filovirus activity and favorable cytotoxicity profiles. Molecular docking indicates these compounds target conserved structural and functional domains—primarily the VP35 and L proteins—which may disrupt essential viral replication and immune antagonism. Furthermore, systematic combinatorial screening revealed three highly synergistic compound pairs, notably NCGC00113249-01 and NCGC00118008-01, demonstrating robust cross-species efficacy. By targeting conserved vulnerabilities across the filovirus family, this integrated *in silico* and *in vitro* pipeline provides a scalable framework to rapidly nominate and optimize synergistic therapeutic regimens against both endemic and emerging viral threats including BDBV.

**Graphical Abstract:** 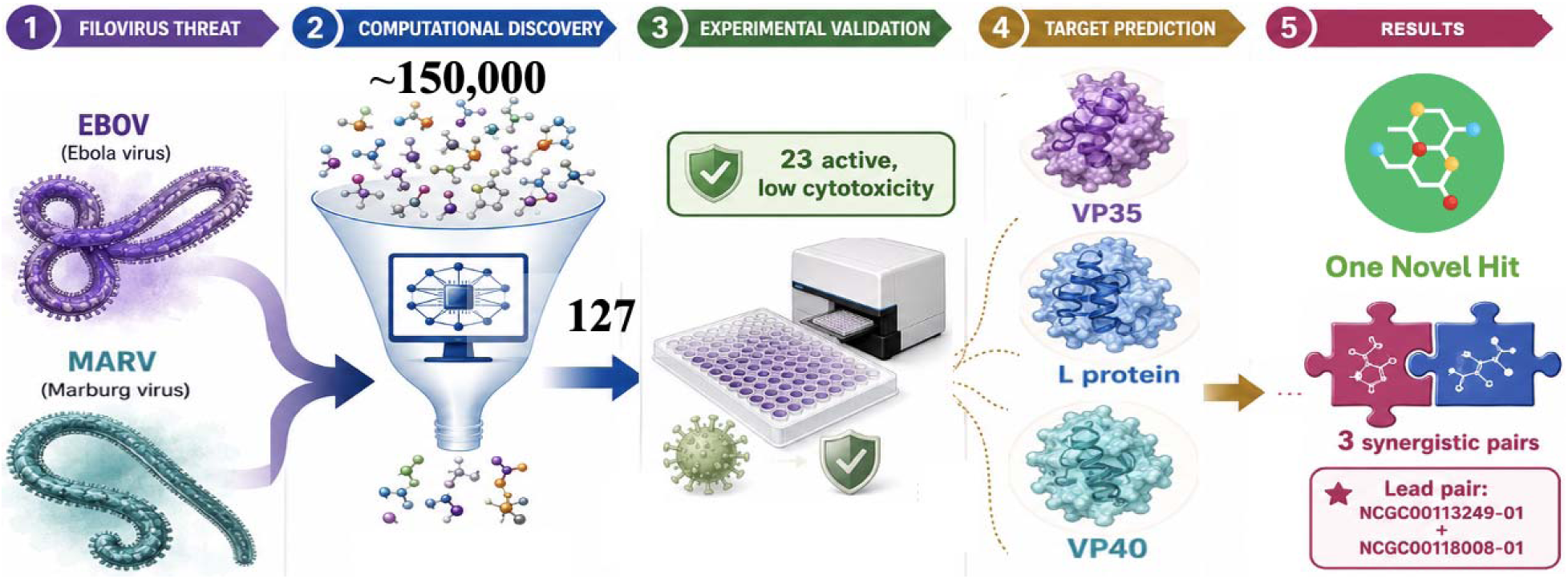

## Introduction

Filoviruses such as Ebola virus (EBOV) and Marburg virus (MARV) are among the National Institute of Allergy and Infectious Diseases’ (NIAID) emerging infectious disease research areas of interest list, and are classified by HHS as tier 1 and category A select agents. Outbreaks of EBOV and MARV are characterized by abrupt onset of disease, person-to-person transmission through direct contact with blood and body fluids, and high case fatality rates that can reach up to 90%.^1^ The 2014–2016 West African EBOV epidemic, which resulted in more than 28,000 cases and over 11,000 deaths, remains the largest filovirus outbreak recorded.^2^ Recent EBOV and MARV outbreaks across Central, East, and West Africa, including events in the Democratic Republic of the Congo, Uganda, Tanzania, Rwanda, Equatorial Guinea, Ghana, and Ethiopia, demonstrate the continued recurrence of filovirus emergence across disperse geographic regions.

The severity of recent Ebola outbreaks underscores that filovirus frequent emergence remains an immediate and escalating global-health threat, not a historical concern, with each new event exposing persistent gaps in rapid therapeutic deployment, outbreak containment, and regional preparedness. The recurrence of EBOV and MARV outbreaks has unfortunately validated our warning that future filovirus epidemics should be expected, not treated as exceptional events, underscoring the need for sustained investment in surveillance, basic virology, epidemiology, and antiviral discovery to reduce their public-health impact.^3^

Therapeutic options for filoviruses remain limited. Although two monoclonal antibody products are licensed for Zaire ebolavirus disease, Inmazeb and Ebanga, and an effective rVSV-ZEBOV vaccine, ERVEBO, is available for prevention of Zaire ebolavirus infection, these countermeasures are species-specific and do not address all clinically important ebolaviruses.^4^ Critically, a current outbreak is caused by Bundibugyo (BDBV) virus, for which there are no approved vaccines or treatments, underscoring the continued need for broad-spectrum filovirus therapeutics.^5^ These countermeasures are virus-specific and do not extend to MARV, for which no approved therapies or vaccines exist.^6^ Even for EBOV, manufacturing capacity and supply-chain limitations have restricted widespread access during outbreaks.^7^ A strong scientific rationale therefore exists for developing pan-filovirus, small molecule inhibitors and/or their combinations. EBOV and MARV share conserved viral proteins and replication mechanisms, including the L protein RNA-directed polymerase (RdRp) and its cofactor VP35, which are essential for viral RNA synthesis and innate immune evasion.^8^ Structural studies have highlighted similarities in these complexes across the filoviruses, suggesting that inhibitors targeting these processes could achieve cross-species activity.^9,10^ Broad-spectrum approaches would enable faster deployment during outbreaks, provide a therapeutic bridge while vaccines or pathogen-specific drugs are mobilized, and reduce dependence on precise strain identification early in an epidemic.^11^

This rationale is especially relevant to Bundibugyo virus (BDBV), the etiologic agent of a current Ebola outbreak, for which no approved vaccine or therapeutic is available. Moreover, existing EBOV-specific therapeutics (Inmazeb and Ebanga monoclonal antibodies, ERVEBO vaccine) has limited efficacy against BDBV because although BDBV exhibits sequence identity values exceeding 72% across six of the seven structural proteins, the only exception – glycoprotein (GP) consistently shows lower conservation across all analyzed filoviridae species. The GP mismatch means antibody-based therapies, targeting exactly this protein, will likely need re-optimization. However, although Bundibugyo virus was not directly tested in this study, the conservation of other essential filovirus proteins and replication mechanisms supports the expectation that inhibitors targeting the most conserved VP35, VP40, and, especially, RdRp of the L protein, may retain activity of both EBOV and MARV. This offers a strategic path to develop broad-spectrum therapeutics battling both current and future emerging pathogens. In addition, the strategy targeting a combination of virus mechanisms that perturb complementary stages of the viral life cycle may be more durable against species-specific differences than single-agent approaches. Therefore, the discovery of inhibitors that target across filovirus activity and work in observed synergy against both EBOV and MARV provide a strong rationale for prioritizing such compounds for follow-up testing against Bundibugyo virus.

Beyond direct clinical benefit, such agents would improve outbreak control and preparedness. Early administration of antivirals can reduce viral load and transmission, thereby limiting epidemic size and mortality.^12^ A single broadly active drug would also simplify stockpiling and distribution, particularly in resource-limited regions. Moreover, broad-spectrum inhibitors align with global health security priorities, addressing WHO’s call for countermeasures against high-risk pathogens and enhancing resilience to future “Disease X” events.^13^

Taken together, the urgent need for effective MARV therapies, the limitations of EBOV-specific interventions, and the conservation of key filovirus replication and assembly mechanisms provide a strong rationale for developing cross-filovirus small-molecule inhibitors and rational antiviral combinations. In this study, we designed an integrated prioritization workflow (**Figure 1**) to move from large-scale computational screening to experimentally validated individual and combination candidates. First, a curated MARV QSAR model built on qHTS data and previously developed EBOV antiviral and cytotoxicity QSAR models were used to prioritize compounds predicted to combine antiviral activity, acceptable cytotoxicity, applicability-domain support, and chemical diversity. Second, selected compounds were tested under BSL-4 conditions to identify experimentally active and non-cytotoxic candidates with activity against authentic MARV and EBOV. Third, conserved-target docking across EBOV and MARV protein structures was used to generate putative mechanism-of-action hypotheses for prioritized hits. Finally, selected compounds were evaluated in 6 × 6 combination matrices to determine whether mechanistically complementary candidates could produce reproducible cross-virus synergy without corresponding cytotoxicity. Thus, the novelty of this work is not only the identification of individual anti-filovirus hits, but the construction of a decision pipeline that links computational enrichment, validation with authentic filoviruses in BSL-4 containment, target-hypothesis generation, and combination prioritization in a single anti-filovirus discovery strategy.

**Figure 1.**
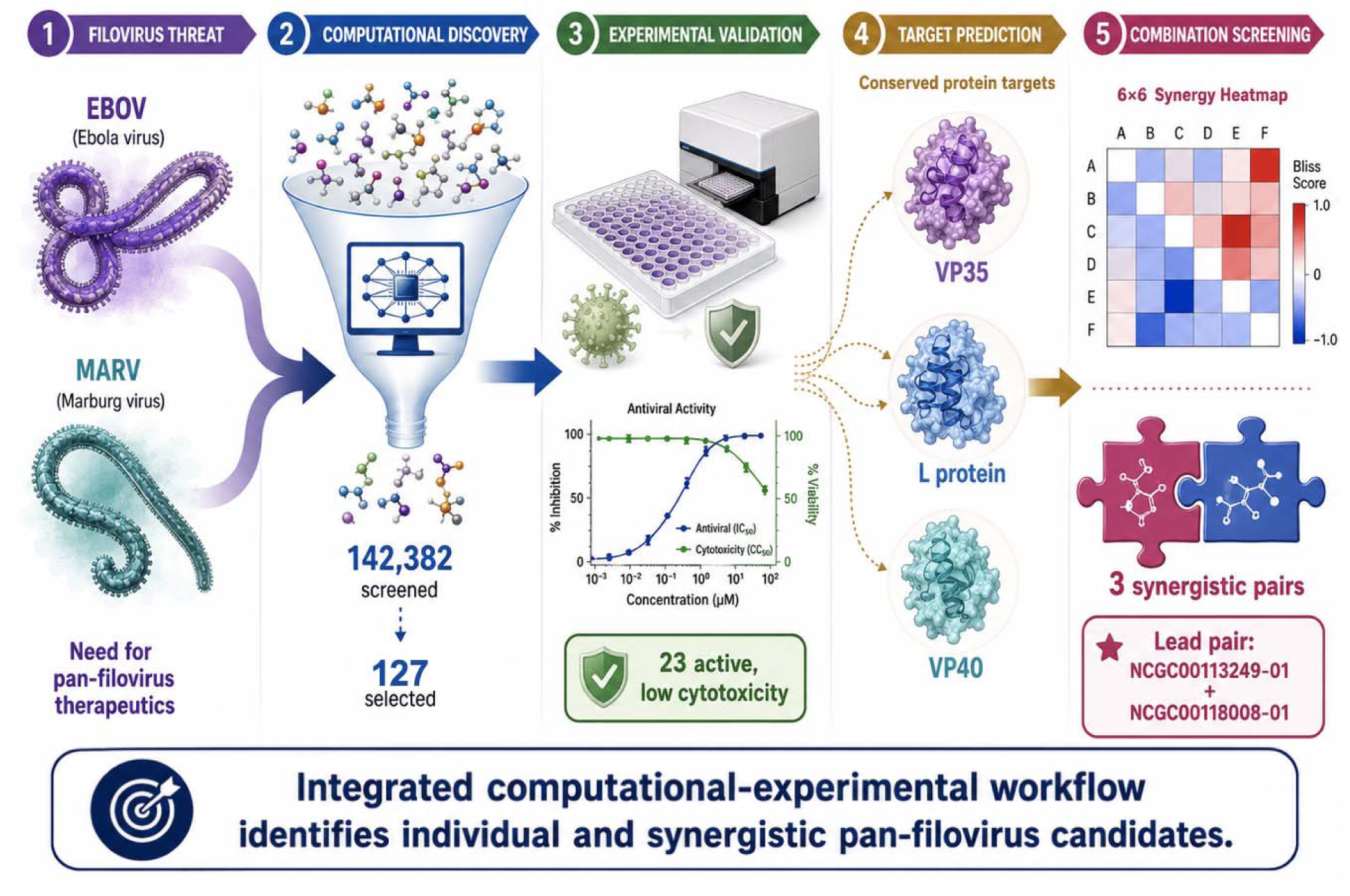
Study Overview

## Methods

### Marburg Virus

#### QSAR Model Data Collection and Curation

For Marburg Virus we collected two assays from PubChem^14^ (PubChem AID: 540276 and 720532) titled “qHTS for inhibitors of binding or entry into cells for Marburg Virus”. The assay data were biologically and chemically curated according to the best practices in the field.^15–17^ These datasets underwent extensive biological data curation due to a large degree of uncertainty in the assay results reported that may directly affect the quality of the models.^18^ Specifically, we only kept data associated with curve class^19^ +/− 1.1, 1.2, 2.1, or 2.2 and set all (+) curve classes to inactive. If a compound had an activity score ≥ 85 in the screen, the compound was classified as “active”.^20^ Similarly, if a compound had an activity score <85 in the screen, the compound was classified as “inactive”. All data points that have >85 activity score but had “excluded data points” (i.e., masked data points) were considered inactive due to the uncertainty in their dose response data. We set binary activity calls for the data using a threshold of >85% activity score to define an active compound. We removed all assay results annotated as “inconclusive”. Any entries with missing activity scores or chemical structures were removed. For each entry, we followed one of the two appropriate procedures for handling duplicates: (i) if the outcomes of all duplicates were concordant, one record was kept with the respective outcome; (ii) if any outcomes disagreed, all were removed. As for chemical curation, we removed mixtures, inorganics, and large organic compounds, removed counterions, cleaned and neutralized salts, and normalized chemotypes using the ChemAxon Standardizer software. The final curated dataset was 5,240 compounds, including 509 actives and 4731 inactives.

#### Model Preparation and Performance

The resulting Marburg dataset was used in KNIME’s RDKit^21^ node to calculate extended-connectivity fingerprints with a radius of 2 (ECFP2) and 2048 bits, as well as Molecular ACCess System (MACCS) Fingerprints for use in modeling. Models were developed using the Random Forrest (RF) algorithm as described previously.^22^ QSAR Models were developed and validated according to the best practices in cheminformatics.^23–25^ In the RF Algorithm, trees were decorrelated via bootstrapping with replacement. Ten rounds of Y-randomization was performed for the model to avoid chance correlations.^26^

For binary classification models, compounds labeled as inhibitors were classified as positive (class 1), and non-inhibitors (or activators) were classified as negative (class 0). We primarily focused on improving the positive predictive value (PPV) rather than other common metrics like balanced accuracy (BA). By focusing on PPV, we ensured that the final model would be better suited for the practical constraints of high-throughput virtual screening.^27^

For Ebola virus models, we used our previously curated antiviral activity and host cytotoxicity datasets, together with their associated QSAR models from our earlier study.^20^ We did not introduce additional data into the training sets, nor did we retrain or recalibrate the models. Instead, we applied the previously validated classification models directly to newly generated chemical datasets using the same descriptor spaces and algorithmic configurations defined during model development.

#### Virtual Screening

Two internal NCATS libraries containing 44,953 and 97,429 compounds, respectively, were used for prediction and prioritization. The 2D structures were processed following the curation protocols detailed above. We utilized an applicability domain (AD) to identify “reliable” and “unreliable” regions of the chemical space for predictions. Compounds were prioritized for experimental testing based on concordance across the MARV and EBOV prediction workflow, applicability-domain status, and chemical diversity. Compounds predicted to have antiviral activity while maintaining acceptable cytotoxicity profiles were preferentially retained. This strategy was intended to enrich for compounds with potential cross-filovirus activity rather than compounds predicted to act against only one virus.

To reduce redundant selection during the virtual screening, compounds were filtered based on chemotypes when selecting the best candidates for *in vitro* testing. No more than five compounds with the same chemotype were selected to ensure chemical diversity. This procedure was used to select 256 compounds from each of the two datasets. Finally, a total of 125 plus two reference compounds were selected for cytotoxicity testing in the Huh-7 cell line.

#### Molecular Docking

Structure-based target prediction was performed via molecular docking using Molegro Virtual Docker (MVD) v7.0.^28^ To ensure a robust assignment of putative viral targets, a consensus-based strategy was employed by conducting docking simulations against homologous protein pairs from EBOV and MARV.

We have focused on the conserved proteins, as more promising targets for creation of polyviral agents within the filoviridae.^29^ Therefore, the following viral proteins were evaluated: nucleoprotein (NP), viral proteins VP24, VP30, VP35, and VP40, glycoprotein (GP), and the large RNA-dependent RNA polymerase (L protein). For targets with experimentally characterized binding sites, docking grids were centered on the reported active regions. In the absence of validated pockets, potential binding cavities were identified using the cavity detection algorithm implemented in MVD.

Experimentally determined structures were retrieved from the Protein Data Bank (PDB).^30^ The following PDB entries were employed: NP: EBOV (4ZTG^31^) and MARV (7F1M^32^); VP24: EBOV (4M0Q^33^) and MARV (4OR8^34^); VP35: EBOV (4IBC^35^) and MARV (4GH9^36^); VP30: EBOV (9JQK^37^) and MARV (5T3W^38^); VP40: EBOV (4LDB^39^) and MARV (5B0V^40^); GP: EBOV (6G95^41^) and MARV (5UQY^42^); L protein: EBOV (8JSL^43^).

Since no experimentally resolved structure was available for the MARV L protein, a three-dimensional model was generated by homology modeling using the SWISS-MODEL server.^44^ The model was built with the ProMod3 engine employing the cryo-EM structure PDB 9IP4 (chain A) as template,^45^ presenting a sequence identity of 94.04% and GMQE ≈ 0.53, indicating high structural reliability for downstream docking analyses. Protein structures were prepared by removing crystallographic water molecules and non-relevant ligands, followed by addition of hydrogen atoms and assignment of appropriate protonation states. Ligand structures were energy-minimized prior to docking.

Docking simulations were performed using Molegro Virtual Docker.^28^ Each compound yielded one final consensus score per viral target. Targets were ranked for every compound based on these final scores, and the top-ranked protein was assigned as the predicted molecular target. This docking-based consensus strategy integrates dual scoring (MolDock and Rerank) with dual-virus structural information (EBOV/MARV), aiming to reduce target-specific bias and improve robustness of target assignment.

#### Mixture Selection

Similar to our previous studies, in order to find the synergistic combinations, we have chosen compounds with different mechanisms of action and/or targeting the virus at different stages of its lifecycle, which increases the probability of synergy between drugs.^46^ A combination of *in silico* tools such as Chemotext,^47^ recently developed IDKG,^48^ and QSAR models of major drug-drug interactions^49^ were used to determine if compounds had been previously tested together and if any negative drug-drug interactions were to be expected. If no information about the mechanism of these compounds was retrieved using text mining, we assigned the putative stage according to docking results described above. In the end, we prioritized 16 combinations of 10 compounds for testing against both viruses. To use drug-drug interaction filter, we calculated simplex descriptors as described elsewhere.^50^

#### Biosafety

All work with infectious virus was performed at CDC by trained personnel in a biosafety level 4 (BSL-4) laboratory. Experiments involving recombinant DNA were approved by the CDC Institutional Biosafety Committee.

#### Cells and Viruses

Huh7 cells (Apath LLC, Brooklyn, NY) were grown in DMEM supplemented with 1x nonessential amino acids (NEAA; ThermoFisher), and 10% (v/v) fetal calf serum (FCS; Cytiva, Marlborough, MA). Infectious reporter viruses expressing ZsGreen (ZsG) were generated as described previously; Marburg (MARV-ZsG; Bat371 Uganda20070 strain Albarino et al. 2013 24074586) and Ebola (EBOV-ZsG; Makona strain Albarino et al. 2015 26122472). Cells were maintained at 37°C with 5% CO_2_. For fluorescent measurements, cells were incubated with FluoroBrite DMEM (ThermoFisher) supplemented with 1x NEAA, 10% FCS, and 1x GlutaMax (ThermoFisher).

#### Determination of Cytotoxicity through Huh-7 Cell Viability Assay

The cytotoxicity testing occurred at the National Institute of Health’s National Center for Advancing Translational Sciences. In the Huh-7 cell line, a compound was considered “non-toxic” if the associated AC50 > 10.0 μM; whereas a compound was considered “toxic” if the AC50 ≤ 10.0 μM or the curve class was 4, indicating no response. Only compounds with dose-response curve classes of 1.1, 1.2, 2.1, 2.2, and 4 were considered.

### Single Point Antiviral Activity Testing

125 compounds were initially screened in single dose of 20-30uM and included positive virus inhibitor remdesivir at 20nM and a cytotoxicity control staurosporine at 120nM. Huh7 cells were seeded at 4000 cells/well in opaque black 384-well plates. Compounds were added the following day using an Echo acoustic liquid handler (Beckman Coulter, Indianapolis, IN), such that the final concentration is 0.2% DMSO. In the BSL-4 lab, MARV-ZsG was added at a multiplicity of infection (MOI) of 0.2 or EBOV-ZsG at a MOI of 0.3 2-hours after compound addition. Three days later, ZsGreen fluorescence activity was measured using a Synergy microplate reader (Agilent BioTek, Santa Clara, CA). Fluorescence values were normalized to uninfected and DMSO-only control wells, and median values from 5 replicates calculated. Viability was assessed in parallel from compound treated uninfected cells using CellTiter-Glo according to manufacturer’s instructions (Promega, Madison, WI). 38 compounds were selected for further testing given greater than 50% virus inhibition and 70% viability.

#### Dose Response Activity

Concentration-response experiments were performed in 384-well plates with four replicates of each compound in an 8-point, half-log serial dilution. Huh7 cells were seeded at a density of 4000 cells/ well. The following day, compounds were added starting at 50µM in 0.5% (v/v) DMSO and incubated 2 hours before virus addition. 72 hours post infection, ZsG fluorescence was measured and infection calculated by normalization to uninfected and DMSO-treated wells. Cell viability was assessed in parallel. Remdesivir (starting 0.5µM) and staurosporine (starting 15µM) were included as positive inhibition and cytotoxicity controls, respectively. For compounds showing inhibition of greater than 50%, the concentration-response dose was calculated using a 4-parameter variable slope logistic model by nonlinear regression to determine the EC_50_ and CC_50_ values (GraphPad Prism v11.01).

#### Matrix Screening of Compound Mixtures

Selected compound pairs were evaluated for combination activity using a 6 × 6 dose–response matrix format in MARV, EBOV, and CellTiter-Glo viability assays. Because single-agent antiviral activity was primarily observed at higher concentrations, compounds were tested at 0, 4, 6, 8, 12, and 20 µM. Each matrix contained all pairwise concentration combinations for the two compounds, allowing antiviral activity to be assessed across a range of single-agent and combined exposures. Duplicate compound combinations were included when available to assess reproducibility of the observed interaction patterns.

Combination effects were analyzed using the Bliss independence model. Deviation from Bliss additivity was calculated for each concentration pair, and negative ΔBliss values were interpreted as enhanced inhibition beyond the expected additive effect. DBSumNeg, defined as the sum of the negative ΔBliss values across the matrix, was used as a summary measure of synergistic activity. In parallel, CTG viability matrices were reviewed to determine whether antiviral effects were associated with compound-induced cytotoxicity. Combinations showing reproducible antiviral synergy in MARV and/or EBOV, without a corresponding pattern of broad toxicity in the CTG assay, were prioritized for further evaluation.

## Results and Discussion

The central goal of this study was to identify anti-filovirus candidates through a staged prioritization strategy rather than through a single screening endpoint. Each step was designed to reduce number of chemicals while increasing biological confidence. Rigorous data curation substantially decreased the dataset size and increased our confidence in modeling results. QSAR-based virtual screening first enriched large NCATS chemical libraries for compounds predicted to have antiviral activity and acceptable cytotoxicity. BSL-4 antiviral testing then distinguished computationally prioritized candidates with measurable activity against MARV and EBOV. Molecular docking of conserved viral targets generated putative mechanistic hypotheses for the active compounds, enabling candidate interpretation in the context of viral entry, replication, immune antagonism, and assembly pathways. Finally, matrix-based combination testing determined whether selected compounds with complementary predicted targets could produce reproducible antiviral synergy across both MARV and EBOV without a proportional increase in cytotoxicity. This workflow reduced 142,382 starting compounds to 127 experimentally tested candidates, 38 dose–response-tested compounds, 23 favorable anti-filovirus hits, 4 prioritized single-agent candidates, and 3 cross-virus synergistic compound pairs. **Table 1** summarizes the decision pipeline used to move from large-scale *in silico* screening to experimentally validated single-agent and combination candidates.

**Table 1.**
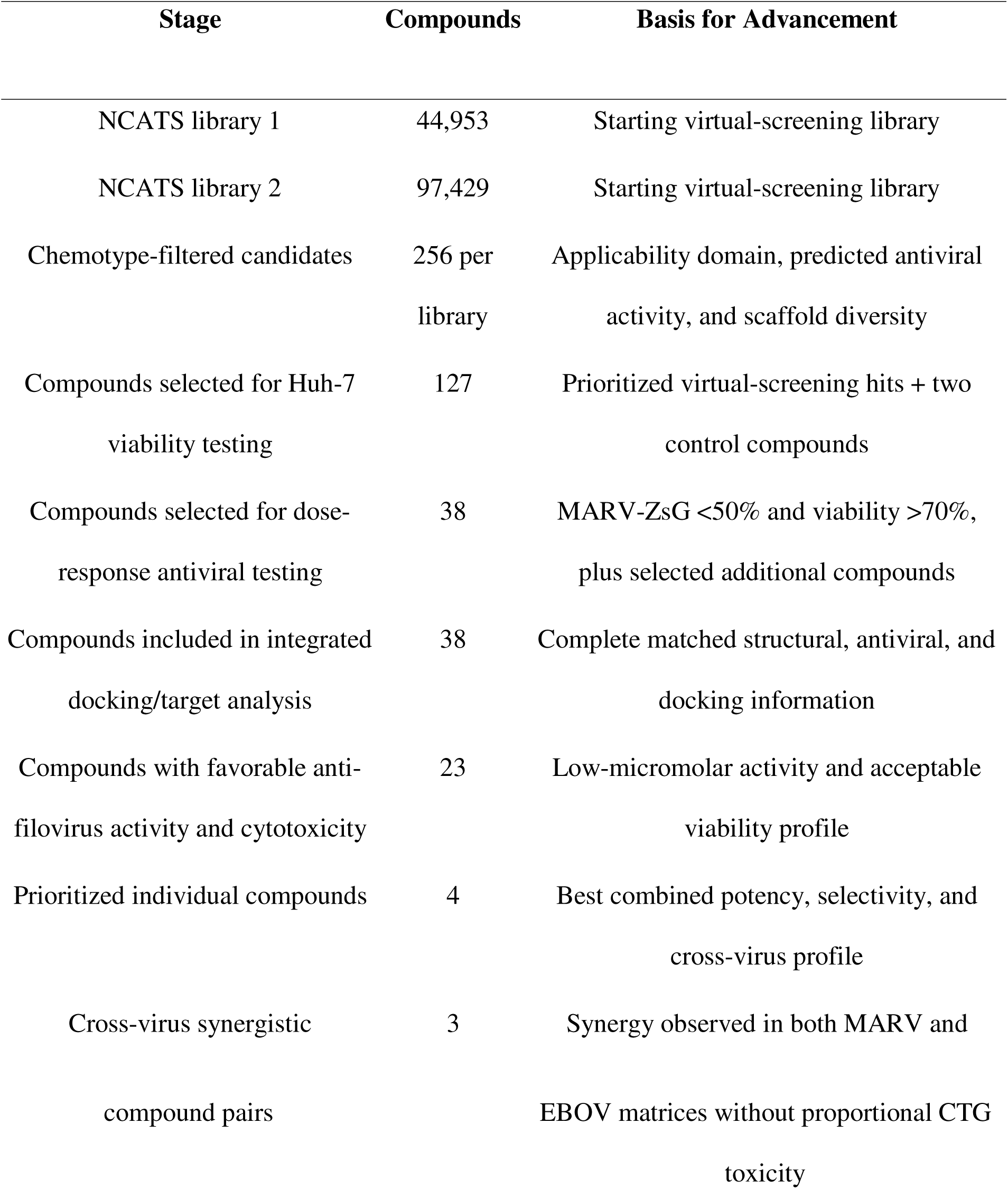
Stepwise prioritization of anti-filovirus candidates from virtual screening to cross-virus synergistic combinations.

### Model Performance, Virtual Screening, and Validation

The Marburg QSAR model built and validated in this study for the MARV data was deemed robust and statistically valid. Our MARV models primarily focused on positive predictive value (PPV) rather than other common metrics to meet the demands of high-throughput virtual screening (HTVS) campaigns, where minimizing false positives is critical. The MARV activity model developed in this study was evaluated using five-fold external cross-validation. At an activity threshold of 85%, the dataset comprises 509 active and 4,731 inactive compounds, corresponding to a baseline active prevalence of approximately 10%. Despite this pronounced class imbalance, the model achieves a positive predictive value (PPV) of 0.50, representing an approximately five-fold enrichment of active compounds relative to random selection. This demonstrates that the model substantially increases the likelihood that predicted actives are true positives, supporting its utility for downstream experimental prioritization under conditions of severe class imbalance. The Ebola QSAR models we used^20^ were previously shown to exhibit robust predictive performance, with five-fold external cross-validation yielding correct classification rates (CCR) above 0.60 across both antiviral activity and host cytotoxicity endpoints, and with no Y-randomized models achieving comparable performance, indicating that model predictions were not driven by chance correlations. The subsequent antiviral validation therefore served as a direct experimental test of whether the QSAR workflow could concentrate biologically active compounds from a much larger chemical space.

QSAR-based virtual screening was used as the first prioritization step to enrich two NCATS compound libraries for candidates with predicted cross-filovirus activity and acceptable cytotoxicity. The combined screening space contained 142,382 compounds, consisting of 44,953 compounds from library 1 and 97,429 compounds from library 2. Compounds were advanced only when they met multiple criteria, including applicability-domain support, predicted antiviral activity, and chemical diversity after chemotype filtering. This strategy reduced the initial screening space to 125 compounds selected for experimental validation, corresponding to less than 0.1% of the starting chemical space. Overall, the virtual screening stage was designed to maximize the probability that experimentally tested compounds would be active, non-cytotoxic, and chemically diverse, rather than capturing every possible active chemotype. The success of this strategy was evaluated experimentally through Huh-7 viability testing, single-point MARV antiviral screening, and subsequent MARV/EBOV dose–response assays.

#### Docking Hit Compounds to Conserved Filovirus Proteins

For each of the selected compounds, the viral protein presenting the highest normalized consensus score, calculated as the arithmetic mean of scaled scores across Ebola virus (EBOV) and Marburg virus (MARV) homologous structures, was defined as the predicted primary target (**Supporting File 1**). Based on this criterion, the docking-based target assignment revealed a distinct distribution, with a significant preference for VP35, L protein, GP, and VP40 (**Table 2**). Specifically, among final 38 hits, the L protein and VP35 were identified as the top-ranked target (12 and 11 compounds, respectively), followed by. the GP (8 compounds), and VP40 (6 compounds) of the total, respectively.

**Table 2.**
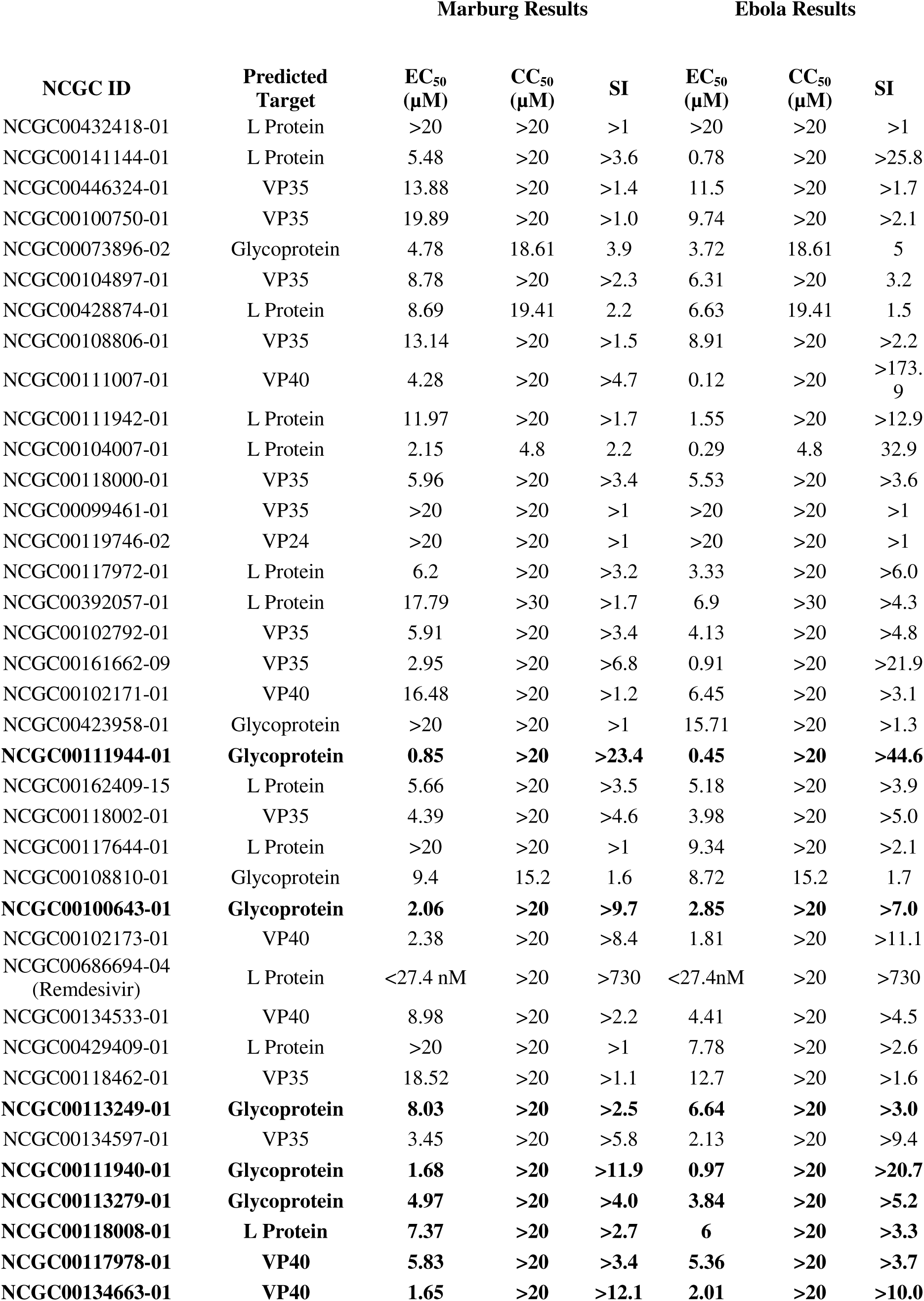
Comparative antiviral activity, cytotoxicity, and predicted viral target class for selected compounds against Marburg and Ebola virus assays. For each compound, the predicted target column was selected using the highest normalized consensus score of molecular docking averaged across EBOV and MARV homologous structures. Compounds that were prioritized are in bold.

The L protein, as the catalytic subunit of the RNA-dependent RNA polymerase (RdRp), emerged as a co-dominant target. Given its indispensable role in viral transcription and genome replication, its prominence among the predicted targets aligns with core antiviral drug discovery strategies. The appearance of VP35 is consistent with its multifaceted role in the viral life cycle, serving as an essential polymerase cofactor and a key antagonist of the host interferon response. The well-characterized RNA-binding domain (RBD) and protein–protein interaction interfaces of VP35 provide druggable cavities that are chemically compatible with diverse small-molecule scaffolds, likely explaining its enrichment in the docking consensus.

One important consideration is that the MARV training data were derived from assays annotated for inhibition of viral binding or entry, whereas the docking analysis predicted several prioritized compounds to interact with internal viral proteins involved in replication or assembly. This apparent distinction may reflect the limitations of assay annotation, the possibility that the original screening readouts captured downstream effects beyond entry, or the ability of chemically related compounds to affect multiple stages of the viral life cycle. Therefore, the docking results should be interpreted as target hypotheses rather than definitive mechanistic assignments.

Within the prioritization framework, docking served two purposes. First, it provided a structure-based rationale for why selected compounds might retain activity across divergent filoviruses by identifying putative interactions with conserved viral proteins. Second, it informed combination selection by assigning active compounds to potentially complementary viral functions. The enrichment of predicted interactions with L and G protein, VP35 and VP40 was particularly relevant because these proteins represent distinct but essential stages of the filovirus life cycle: viral entry, polymerase cofactor function and innate immune antagonism, viral RNA synthesis, and virion assembly/budding. Therefore, while the docking results do not establish direct target engagement, they provide a mechanistic framework for interpreting both single-agent cross-virus activity and the observed synergy among selected compound pairs.

#### Antiviral Activity Assay

The dose–response results are summarized in Table 2 and reported fully in Supporting File 2. Several compounds showed single-digit micromolar EC_50_ values against one or both viruses, with CC_50_ values above the highest tested concentration and corresponding selectivity indices consistent with a favorable preliminary antiviral profile. Remdesivir, included as a benchmark antiviral control, produced the strongest apparent activity, with EC50 values below 27.4 nM and SI values greater than 730 in both assays. Importantly, multiple non-control compounds retained low-micromolar activity against both MARV and EBOV, supporting the ability of the workflow to identify cross-virus active candidates. In addition to these, we also used Staurosporine (NCGC00162400-01) as a negative control to benchmark the CC_50_ results **(Supporting File 2**).

The most compelling single-agent candidates were those that combined three properties: measurable activity against both MARV and EBOV, acceptable selectivity indices, and a putative target assignment involving a conserved viral function. This integrated interpretation prioritized compounds not solely by potency, but by their combined cross-virus activity, selectivity, and mechanistic plausibility. Compounds assigned to L protein, VP35, VP40, and glycoprotein included several of the strongest candidates, supporting the concept that conserved filovirus proteins can provide useful organizing targets for broad-spectrum antiviral discovery. These results justify advancing selected compounds into orthogonal antiviral validation, target-engagement studies, and medicinal chemistry optimization.

#### Prioritization of Cross-Filovirus Single-Agent Candidates

Based on combined potency, selectivity, cross-virus activity, and target-hypothesis support, compounds **NCGC00111944-01**, **NCGC00100643-01**, **NCGC00111940-01**, and **NCGC00134663-01** were prioritized as the strongest non-control single-agent candidates for follow-up evaluation. To avoid prioritizing compounds based on potency alone, single-agent candidates were evaluated using an integrated profile that considered MARV activity, EBOV activity, cytotoxicity, selectivity index, and predicted target class. This approach identified the strongest candidates as those with reproducible activity across both viral assays and no evidence of broad cytotoxicity at active concentrations.

This prioritization strategy also helped distinguish benchmark-like highly potent compounds from structurally and mechanistically informative candidates. For example, Remdesivir (NCGC00686694-04) was our control compound and showed the strongest apparent potency and selectivity across both viruses, whereas additional compounds with low-micromolar activity and acceptable SI values provided chemically diverse candidates for follow-up optimization. The target-prediction results further strengthened this prioritization by linking active compounds to conserved viral processes, including L protein-mediated RNA synthesis, VP35-associated replication-complex and immune-antagonist functions, VP40-mediated assembly/budding, and glycoprotein-associated entry. Thus, the single-agent results nominate not only individual compounds but also conserved viral processes that may be exploited for broader anti-filovirus drug discovery.

#### Compound Mixtures

To evaluate whether selected anti-filovirus hits could produce enhanced antiviral activity in combination, we tested compound pairs in 6 × 6 dose–response matrices against MARV and EBOV, with a parallel CellTiter-Glo viability/toxicity assay. The combination matrices were performed across 0, 4, 6, 8, 12, and 20 µM. Synergy was assessed using Bliss independence, with negative ΔBliss values indicating greater inhibition than expected from the additive single-agent response. DBSumNeg, defined as the sum of negative deviations from the Bliss model, was used as the primary summary metric for synergistic antiviral activity.

Combination testing was used as the final prioritization step to determine whether selected single-agent hits could be converted into more effective cross-virus antiviral strategies (**Supporting File 3).** The principal combination candidates included NCGC00113249-01, NCGC00118008-01, and NCGC00117978-01, which showed measurable activity against both viruses and were computationally assigned to GP (viral entry), L protein (replication), and VP40 (egress), respectively. Because the selected compounds were computationally assigned to different conserved viral targets, the matrix studies tested the hypothesis that simultaneous perturbation of complementary filovirus functions on the different stages of viral lifecycle could produce enhanced antiviral activity. The most important finding was that three compound pairs showed concordant synergy across both the MARV and EBOV assays, indicating that these interactions were not restricted to a single filovirus background (**Figure 2; Supporting File 4**).

**Figure 2.**
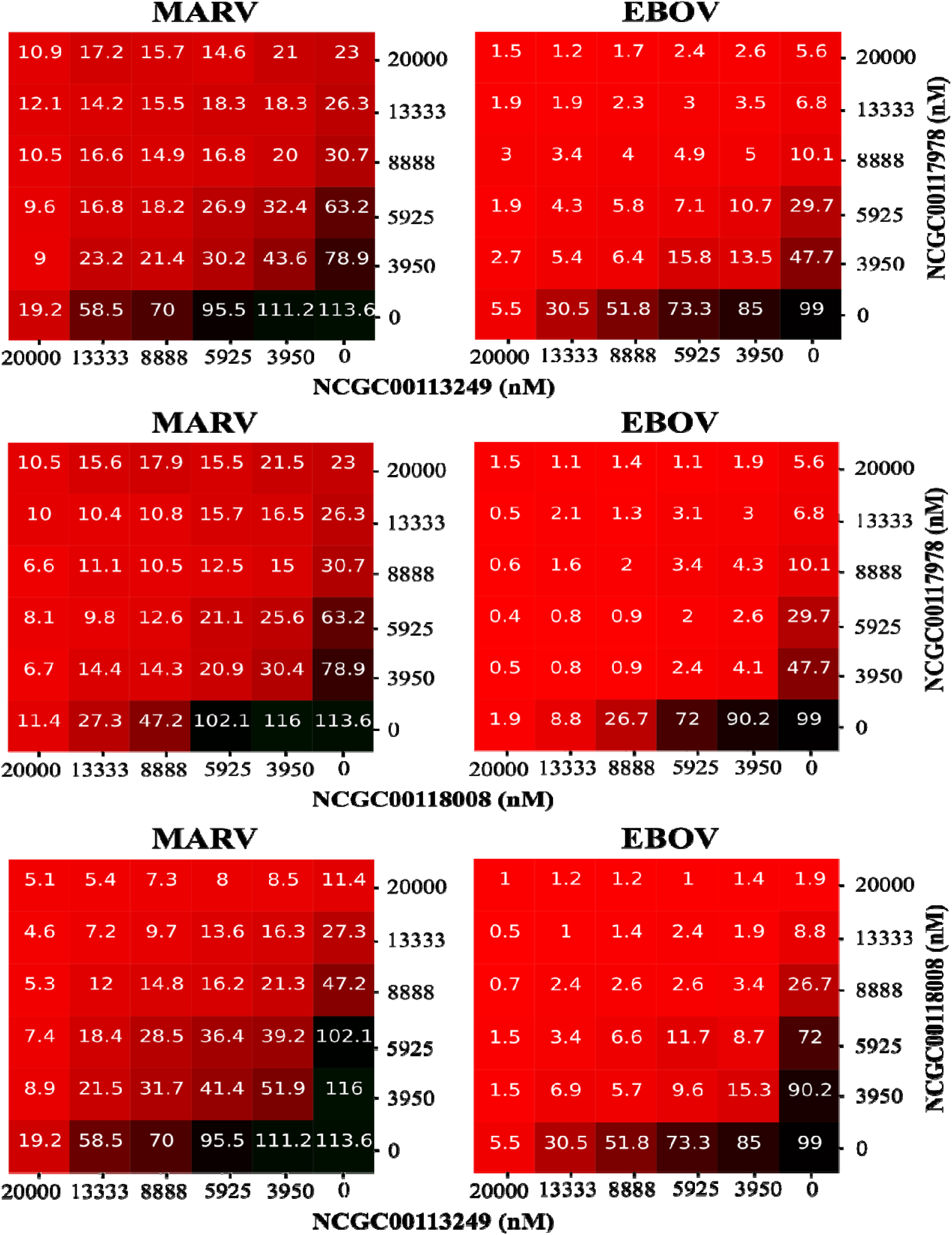
Cross-virus synergistic compound combinations identified in MARV and EBOV matrix assays.

The strongest and most reproducible cross-virus interaction was observed for NCGC00113249-01 + NCGC00118008-01. This pair showed the most favorable Bliss synergy profile across MARV and also ranked as the strongest EBOV synergistic pair. The interaction is mechanistically plausible because NCGC00113249-01 was computationally assigned to GP, whereas NCGC00118008-01 was assigned to the L protein. Our results suggest that combined inhibition of both glycoprotein responsible for viral entry and RdRp responsible for viral replication may enhance antiviral efficacy.

Two additional compound pairs, NCGC00113249-01 + NCGC00117978-01 and NCGC00118008-01 + NCGC00117978-01, also showed cross-virus synergy. NCGC00117978-01 was assigned to VP40, providing a complementary assembly/budding-associated target hypothesis. Together, these three combinations suggest that the most productive antiviral interactions may arise from pairing compounds predicted to affect distinct conserved stages of the filovirus life cycle, including replication-complex function and virion assembly.

EBOV and BDBV are closely related members of the Ebolavirus genus, sharing overall genome organization, homologous viral proteins, and a conserved replication strategy. Although they are distinct species, one of the most biologically important differences is the glycoprotein, which can influence viral entry, tropism, and sensitivity to GP-directed antibodies or inhibitors. Because the remaining viral lifecycle machinery is broadly conserved, we prioritized compound combinations predicted to act outside of GP-dependent entry and instead target complementary conserved stages of infection. On this basis, we suggest NCGC00118008-01 + NCGC00117978-01 as the strongest synergistic pair, because as stated above, NCGC00118008-01 was assigned to the viral L protein, supporting a replication/transcription-associated target hypothesis, while NCGC00117978-01 was assigned to VP40, providing a complementary assembly/budding-associated mechanism. Together, this pair offers mechanistic coverage across distinct and conserved stages of the filovirus lifecycle, supporting its prioritization for activity across both EBOV and BDBV.

The parallel CTG matrices (**Supporting Figure 1**) were essential for interpreting these effects. The lead combinations did not show a proportional pattern of broad cytotoxicity that mirrored the antiviral synergy matrices, suggesting that the observed enhancement was not driven solely by compound-induced loss of cell viability. Although additional cytotoxicity profiling is needed, these data support the prioritization of NCGC00113249-01 + NCGC00118008-01 as the lead cross-virus combination (whole filoviridae except BDBV) and nominate GP/L protein co-perturbation as a testable mechanistic hypothesis for future studies.

Chemical structures are shown for the eight candidate compounds identified (**Table 3**). Compounds **NCGC00111944-01**, **NCGC00100643-01**, **NCGC00113249-01**, **NCGC00111940-01**, **NCGC00113279-01**, and **NCGC00118008-01** were previously known to show MARV activity in qHTS assays for inhibitors of Marburg virus binding or entry (PubChem AID 540276)^51^ and are reported here as newly recognized EBOV-active candidates. **NCGC00134663-01** was identified as a new, potent, and selective anti-filovirus candidate. This prioritized hit set bridged historical MARV activity from a lower-resolution PubChem qHTS binding/entry assay with newly generated live-virus BSL-4 antiviral testing,^51^ revealing compounds with newly observed EBOV activity and identifying one non-control compound with a favorable cross-virus activity and selectivity profile. Importantly, the MARV strain assayed in this study was MARV-ZsG based on 371Bat2007, whereas the historical assay evaluated a VSV pseudotype expressing the MARV GP from the Musoke strain. We are confident in the transferability of results between these systems because 371Bat2007 is a naturally derived MARV isolate and the recombinant ZsG virus preserves the MARV genomic backbone and core viral lifecycle while incorporating a fluorescent reporter to facilitate quantitative readout. The proteins and processes most relevant to broad antiviral activity, including viral entry, polymerase-driven replication/transcription, nucleocapsid function, budding, and host-factor dependence, are highly conserved within MARV relative to the larger divergence observed between MARV and Ravn virus. Thus, although strain-specific differences cannot be excluded, MARV-ZsG/371Bat2007 provides a biologically relevant and more definitive infectious-virus assay system for confirming MARV activity observed in earlier wild-type MARV screening. To further support this assumption, we assessed concordance between the historical MARV assay and our current live-virus results. We retained 15 compounds that were reported active in the historical MARV assay within the broader set of 125 candidates and 12 advanced into the final prioritized set of 38 compounds, and five remained in our list of top prioritized compounds, demonstrating previously unknown high activity and selectivity against EBOV. This assay agreement is notable because cross-assay reproducibility is not always achievable across different assay formats, viral strains, and readout systems. The recovery of selective compounds across both the historical and current assay contexts therefore strengthens confidence that these hits reflect reproducible MARV-relevant antiviral activity rather than assay-specific signal. We additionally labeled the top compounds that showed synergy and were prioritized for combination testing based on their antiviral profile and potential utility in synergistic compound-pair strategies. We also approved each compounds drug-likeness using the STOPLIGHT web portal to score and predict molecular properties, assay liabilities, and pharmacokinetic properties (**Supporting File 5**).^52^

**Table 3.** Prioritized anti-filovirus hit compounds selected for follow-up evaluation including their individual MARV and EBOV assay results. Single compounds are reported in panel A. Panel B reports compounds prioritized in synergetic combinations with the result shown for the highest synergy reported.

| NCGC ID | NCGC00111944-01 | NCGC00100643-01 | NCGC00111940-01 | NCGC00134663-01 |
| --- | --- | --- | --- | --- |
| MARV EC50 (uM) | 0.85 | 2.06 | 1.68 | 1.65 |
| MARV CC50 (uM) | >20 | >20 | >20 | >20 |
| MARV SI | >23.4 | >9.7 | >11.9 | >12.1 |
| EBOV EC50 (uM) | 0.45 | 2.85 | 0.97 | 2.01 |
| EBOV CC50 (uM) | >20 | >20 | >20 | >20 |
| EBOV SI | >44.6 | >7 | >5.2 | >10 |

| NCGC ID | NCGC00113249-01 | NCGC00118008-01 | NCGC00117978-01 | NCGC00113279-01 |
| --- | --- | --- | --- | --- |
| MARV EC50 (uM) | 8.03 | 7.37 | 5.83 | 4.97 |
| MARV CC50 (uM) | >20 | >20 | >20 | >20 |
| MARV SI | >2.5 | >2.7 | >3.4 | >4 |
| EBOV EC50 (uM) | 6.64 | 6 | 5.36 | 3.85 |
| EBOV CC50 (uM) | >20 | >20 | >20 | >20 |
| EBOV SI | >3 | >3.3 | >3.7 | >5.2 |
| Top Synergy with (-DBSum): | NCGC00118008-01<br>MARV: -5.44 | NCGC00113249-01<br>EBOV: -3.97 | NCGC00113249-01<br>MARV: -3.16 | NCGC00117978-01<br>MARV: -0.62 |

Collectively, these findings support the central premise of the study: computational screening, BSL-4 validation, conserved-target assignment by docking, and matrix-based combination testing can be combined into a practical prioritization framework for anti-filovirus discovery. Rather than functioning as independent analyses, each layer increased confidence in the candidates advanced to the next stage. This progression from computational enrichment to biological validation and combination prioritization represents the major conceptual advance of the study.

Several limitations should be considered when interpreting these findings. First, the docking-based target assignments are computational hypotheses and should not be interpreted as direct evidence of target engagement. Biochemical binding assays, viral minigenome assays, VP35 functional assays, VP40 budding assays, or time-of-addition studies will be required to confirm the proposed mechanisms. Second, although the CTG matrices did not indicate that cross-virus synergy was explained by broad cytotoxicity, additional orthogonal cytotoxicity and cell-health assays are needed to define the therapeutic window more rigorously. Third, antiviral validation was performed in Huh-7 cells, and future studies should determine whether the activity profiles extend to additional relevant cell types. Fourth, the current combination studies used a defined concentration range selected to capture single-agent and combination activity; expanded dose-ratio studies will be needed to optimize the most favorable concentration windows. Finally, because the present work focused on MARV and EBOV, additional testing against other filoviruses, including Ravn virus and other ebolaviruses including the currently circulating Bundibugyo virus, will be important to determine and verify the true breadth of activity.

## Conclusions

The ongoing Bundibugyo Ebola outbreak in the Democratic Republic of the Congo and Uganda underscores the critical vulnerability in global health preparedness: the absence of broadly active filovirus countermeasures. To address this translational gap, we have established a highly integrated, generalizable computational and experimental framework that accelerates the discovery of cross-reactive antivirals. By coupling QSAR-guided *in silico* screening and applicability-domain filtering with rigorous BSL-4 phenotypic validation, we successfully distilled a library of over 140,000 compounds into 23 potent, low-micromolar pan-filovirus inhibitors. While empirical target engagement studies remain necessary, molecular docking indicates that these candidates exploit highly conserved structural vulnerabilities across the filovirus proteome—specifically the G, L, VP35, and VP40 proteins. This suggests a multi-nodal disruption of viral entry, replication-complex assembly, immune evasion, and virion budding.

Crucially, our pipeline facilitates the rational design of synergistic therapeutic regimens. Matrix-based combination profiling demonstrated that targeting complementary viral functions yields enhanced efficacy, highlighted by the pairing of NCGC00113249-01 and NCGC00118008-01. Predicted to dually inhibit GP-mediated entry and L protein-driven replication, this lead combination achieved potent synergy without exacerbating cytotoxicity. These data establish a proof-of-concept that mechanistically complementary, computationally prioritized combinations can effectively overcome the limitations of monotherapy in severe viral infections.

Ultimately, this platform provides a scalable blueprint for rapid antiviral drug discovery. Because our lead candidates and synergistic pairs target core, conserved mechanisms of filovirus pathogenesis, they possess a high probability of retaining robust activity against related species, including the currently endemic Bundibugyo virus. Given the urgent clinical need, immediate evaluation of these compounds against emerging Bundibugyo virus species is warranted. Moving forward, this pipeline will guide medicinal chemistry optimization, definitive target-engagement validation, and *in vivo* efficacy profiling to rapidly advance these broad-spectrum synergistic combinations toward clinical utility.

## Acknowledgments

The authors are thankful to Kelli Wilson and Crystal McKnight (both NCATS) for their help with executing the project. This project has been funded in part with Federal funds from the National Institute of Allergy and Infectious Diseases, National Institutes of Health, Department of Health and Human Services, under Contract No. 75N93024C00038; Conselho Nacional de Desenvolvimento Científico e Tecnológico (CNPq, Brazil, grant number 423264/2025-7) awarded as a Pesquisador Visitante Especial (PVE) fellowship/program; Intramural Research Program of the National Institutes of Health (NIH). The contributions of the NIH author(s) were made as part of their official duties as NIH federal employees, are in compliance with agency policy requirements, and are considered Works of the United States Government. However, the findings and conclusions presented in this paper are those of the author(s) and do not necessarily reflect the views of the NIH or the U.S. Department of Health and Human Services. This research was supported in part by CDC core Emerging Infections funding. The findings and conclusions in this report are those of the authors and do not necessarily represent the official position of the Centers for Disease Control and Prevention.

