## Supplementary figures and images for "AI-Driven Discovery and BSL-4 Validation of Cross-Filovirus Ebola-Marburg Inhibitors and their Synergistic Combinations"

### SupportingFigure1-CTG.png

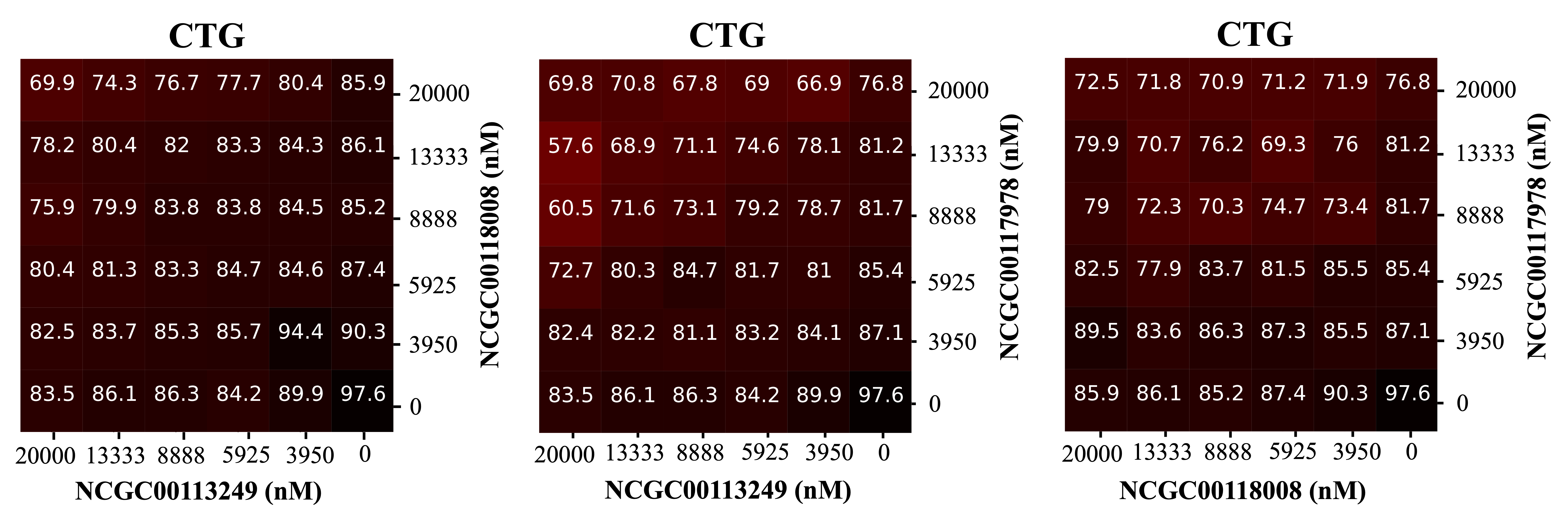
